# Long-lived rodents reveal signatures of positive selection in genes associated with lifespan and eusociality

**DOI:** 10.1101/191999

**Authors:** Arne Sahm, Martin Bens, Karol Szafranski, Susanne Holtze, Marco Groth, Matthias Görlach, Cornelis Calkhoven, Christine Müller, Matthias Schwab, Hans A. Kestler, Alessandro Cellerino, Hynek Burda, Thomas Hildebrandt, Philip Dammann, Matthias Platzer

**Affiliations:** Leibniz Institute on Aging – Fritz Lipmann Institute, Jena, Germany; Department of Reproduction Management, Leibniz Institute for Zoo and Wildlife Research, Berlin, Germany; European Research Institute for the Biology of Ageing, University of Groningen, University Medical Centre Groningen Groningen, The Netherlands; Department of Neurology; Jena University Hospital-Friedrich Schiller University, Jena, Germany; Institute of Medical Systems Biology, Ulm University, Ulm, Germany; Laboratory of Biology Bio@SNS, Scuola Normale Superiore, Pisa, Italy; Department of General Zoology, Faculty of Biology, University of Duisburg-Essen, Essen, Germany; University Hospital, University of Duisburg-Essen, Essen, Germany

## Abstract

The genetic mechanisms that determine lifespan are poorly understood. Most research has been done on short lived animals and it is unclear if these insights can be transferred to long-lived mammals like humans. Some African mole-rats (Bathyergidae) have life expectancies that are multiple times higher than similar sized and phylogenetically closely related rodents. We obtained genomic and transcriptomic data from 17 rodent species and systematically scanned eleven lineages associated with the evolution of longevity and eusociality for positively selected genes (PSGs). The set of 319 PSGs contains regulators of mTOR and is enriched in functional terms associated with (i) processes that are regulated by the mTOR pathway, e.g. translation, autophagy and mitochondrial biogenesis, (ii) the immune system and (iii) antioxidant defense. Analyzing gene expression of PSGs during aging in the long-lived naked mole-rat and up-regulation in the short-lived rat, we found a pattern fitting the antagonistic pleiotropy theory of aging.

## Introduction

Most of the available information about the genetic mechanisms that govern lifespan and aging were obtained by studying single-gene mutations in invertebrates or short-lived, highly inbred vertebrate species. However, it is not clear whether insights about aging relevant genes and pathways gained from these species can be applied to long-lived species like human ^1^. In addition, lifespan extensions under artificial laboratory conditions resulting from single gene mutations or other genetic, pharmacologic and/or lifestyle interventions are far smaller than natural variation of lifespan among species shaped by natural selection. Moreover, it is not clear to what extent genetic variation is responsible for intraspecific heritable differences in lifespan overlaps with the genetic architecture of lifespan macroevolution. As a case in point, maximum lifespan in captivity varies about two orders of magnitude and is positively correlated with body mass in vertebrates ^2,3^, but the two traits are negatively correlated within species, the most extreme example being dog breeds ^4^. Therefore, comparative evolutionary approaches that search for genetic differences between closely related species that are short- and long-lived with respect to their body mass may reveal novel candidate genes and pathways or open new perspectives on known ones.

Rodents are an ideal taxon for such an approach. While the majority of species is short-lived, such as mice, rats and hamsters, there are long-lived exceptions, such as chinchillas, blind mole rats (BMR, *Spalax* sp.) and several African mole-rat species including the naked mole-rat (NMR, *Heterocephalus glaber*) ^5,6^. Furthermore, genome and transcriptome sequences of short- and long-lived species are available and can be used for comparative analysis.

African mole-rats (family Bathyergidae) are subterranean rodents that feed from roots and tubers. The family comprises six genera; for five out of these, maximum lifespan records are available for at least one species. Notably, and in contrast to most other rodents, none of these species has a maximum lifespan of below ten years or below the predictions of the power-law that describes body mass/lifespan relationships in mammals ^6^. At the extreme of this distribution, Zambian mole-rats from the *Fukomys micklemi* clade ^7^ (the best studied representative being the Ansell’s mole-rat *F. anselli*, AMR), the giant mole-rat (GMR, *Fukomys mechowii*) and NMR, have maximum lifespans of at least ca. 20, 22 and 31 years, respectively. These values are 212%, 194% and 368% with respect to the predicted lifespan based on their body mass (^5^, GMR percentage calculated with own lifespan data and same formula). In contrast, the established biomedical model organisms rat (*Rattus norvegicus*) and mouse (*Mus musculus*) have a maximum lifespan of 3.8 and 4 years, respectively, which is 32% and 51% of the predicted value. Remarkably, the greater cane rat (*Thryonomys swinderianus*) that is closely related to the African mole-rats reaches only 28% of the predicted maximum lifespan ^5^ (Fig. 1).

**Figure 1.**
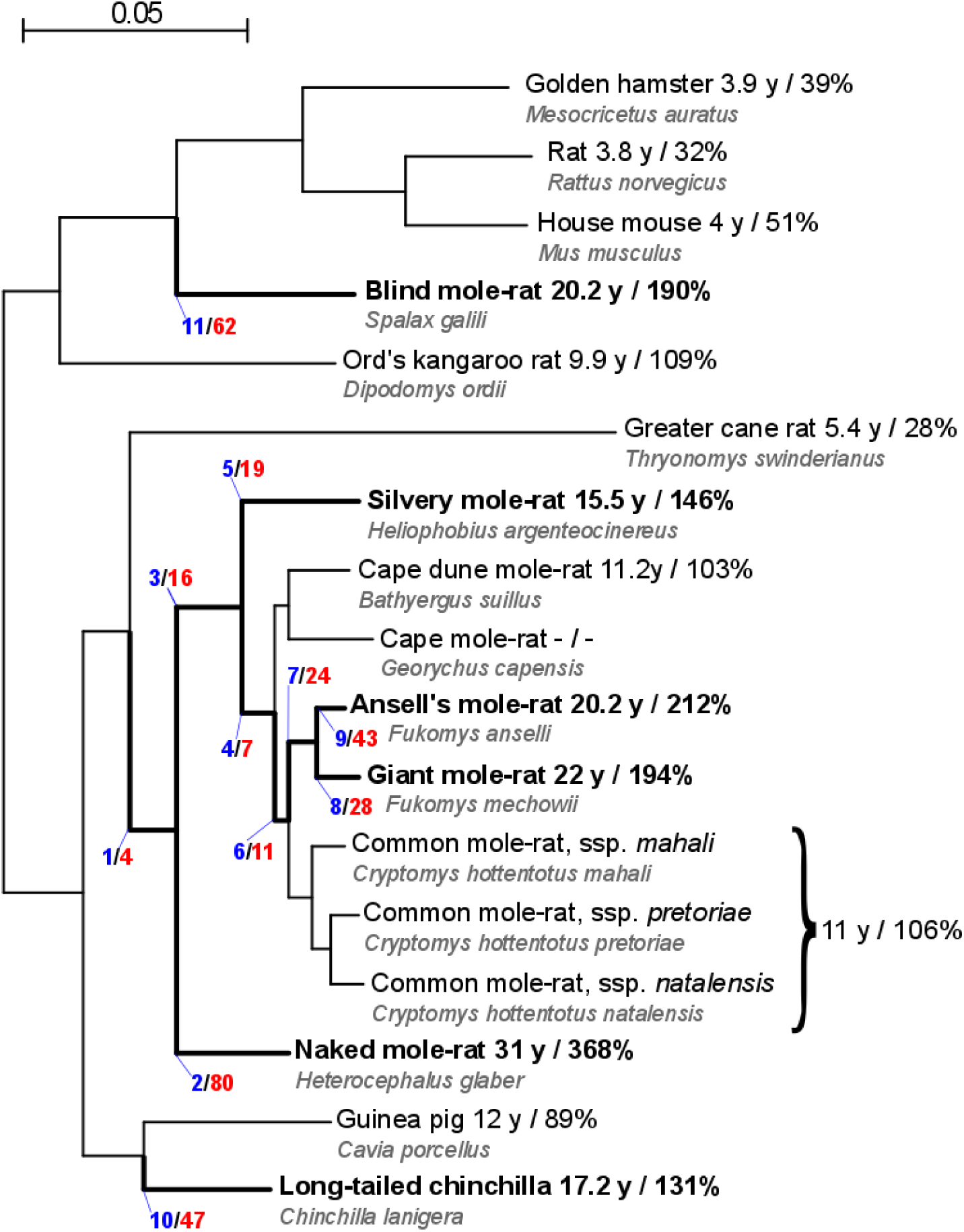
Nucleotide-based phylogeny of the analyzed rodents. Species or branches regarded in the present analyses as long-lived or leading to longevity, respectively, are depicted in bold. The branch numbers used in the text are shown in blue. The numbers of genes with signs of positive selection on the branches are colored in red. The first number after the species name shows the recorded maximum lifespan and the second number is the percentage of the observed vs. expected maximum lifespan based on the respective body mass. The maximum lifespans and ratios were taken from ^5^, except for silvery mole-rat (personal communication by R. Sumbera) and giant mole rat (own data). For these two species, the expected maximum lifespans were calculated with the same mammalian allometric equation used by ^5^. The scale bar represents 0.05 substitutions per site.

Due to a number of unique phenotypes, the NMR became the focus of intensive research ^8^. It was the first vertebrate for which eusociality was discovered (^9^). The NMR shows (i) the longest lifespan among rodents, (ii) no aging-related decline in reproductive and physiological parameters, as well as (iii) no observable aging-related increase in mortality rate ^10^. Among thousands of examined animals only six recently discovered cases of spontaneous tumors have been described ^11,12^. Interestingly, cancer resistance is shared with BMR, which is also long-lived but, despite its name, rather distantly related to African mole-rats (Fig. 1). However, different mechanisms are proposed for cancer resistance in these two taxa. While high-mass hyaluronan mediated early contact inhibition was suggested as a key player in NMR ^13^, a concerted necrotic cell death mechanism in response to hyperproliferation was proposed for BMR ^14^.

The search for signatures of positive selection represents a powerful approach to identify the genetic basis of these unique biological features. Positive selection is the fixation of an allele in a taxon driven by its positive effect on fitness. Once an adaptive phenotype evolved in a given species or evolutionary clade, some of the genes under positive selection likely play a role in it. In protein-coding sequences (CDSs), positive selection results in an increased rate of non-synonymous substitutions as compared to genetic drift. Statistical models based on the ratio of non-synonymous to synonymous substitution rates (*d*_*N*_/*d*_*S*_) are widely used in comparative genomics and allow the identification of specific amino acids within a given gene that changed due to positive selection ^15-17^

Consequently, several studies performed genome-scale scans for positively selected genes (PSGs) in African mole-rats and BMR. The first study ^18^ searched for PSGs on the very long NMR branch in a four-species ^19^ comparison with human as an outgroup and the mouse and rat as further rodents. Among the 142 identified PSG candidates, three were members of a five-protein complex involved in alternative lengthening of the telomeres. The second study ^20^, used ten species with the guinea pig (*Cavia porcellus*) as most closely related species and scanned for PSGs along the branches leading to NMR, Damaraland mole-rat (*Fukomys damarensis*) and their last common ancestor (LCA), identifying 334, 179 and 82 candidates, respectively, including candidates associated with neurotransmission of pain in the NMR. A third study ^21^ used species from all six African mole-rat genera and searched the branch of the LCA of all African mole-rats that follows divergence from the guinea pig. Signs of positive selection were identified in 513 genes, including loci associated with tumorigenesis, aging, morphological development and sociality. All three studies suffer from a methodological limitation that is common in positive selection studies: in none of these, a closer related species than guinea pig was included. As guinea pig is not the closest relative of African mole rats not expressing the phenotypes of interest, it cannot be excluded that fixation of detected signs of positive selection predates – and therefore could not contribute to – the evolution of these phenotypes ^22^. A fourth study ^23^ examined the BMR branch using the Chinese hamster (*Cricetulus griseus*) as the most closely related outgroup. Among the 48 PSG candidates, several were linked to necrosis, inflammation and cancer.

To better resolve the above-mentioned ambiguities and to achieve a higher resolution of positive selection along rodent phylogenetic branches leading to longevity and eusociality, we analyzed genomic and transcriptomic data of 17 species – data from public sources and original data generated for this study. In particular, we generated genomic data for the greater cane rat as a key species absent from previous analysis and for the silvery mole-rat (SMR, *Heliophobius argenteocinereus*). We systematically scanned 11 evolutionary branches (6 corresponding to extant species and 5 to ancestral branches). This approach enables us to date precisely the occurrence of signatures of positive selection with respect to the evolution of the phenotypes of interest on multiple evolutionary branches of rodents. In addition, we generated RNA-seq data from young and old NMRs and laboratory rats (*Rattus norvegicus*) to analyze the overlap between PSGs and genes regulated during aging. Based on this, we discuss the implications of these results on our understanding of the genetic basis of aging, lifespan and sociality.

## Results and Discussion

As starting points for our analysis, we generated CDS libraries for five rodent species (NMR, AMR, GMR, SMR and greater cane rat) based on transcriptomic and genomic data. (Table S1/S2). Together with publicly available rodent CDS catalogs (Table S1), we obtained data for 17 species, including several additional African mole-rats, the chinchilla, BMR and short-lived outgroups like the guinea pig, mouse and rat (Fig. 1). From these sequences, we predicted orthologs and best matching isoforms between the species, calculated alignments and applied multiple times the branch-site test of positive selection ^24^.

Based on the lifespans of the extant species, we regarded six extant as well as five ancestral branches as leading to enhanced longevity and examined them for positive selection (Fig. 1). In total, we detected 341 PSGs (p<0.05, branch-site test). Our PSG assignment is based on nominal p-values, a common approach in genome-wide scans ^21,25,26^ since the main error source of such analyses are alignment errors ^27^ which result in extremely small p-values and therefore cannot be controlled by multiple test corrections. Furthermore, simulations have shown that the empirical false positive rate is very low if an appropriate filtering is used to remove alignment errors and unreliable results ^28^.

Twenty genes were found on multiple branches (Table S3), resulting in a non-redundant set of 319 PSGs (Table S4-S15). Signs of positive selection for the same gene on multiple branches indicate possible parallel evolution. Among those, we found *AMHR2* (anti-Mullerian hormone receptor type 2) to be positively selected both on branch 2 (NMR) and branch 11 (BMR). While AMHR2 plays a role in male fetal development and in ovarian follicle development of the adult female ^29^, no function with regard to aging is described yet. However, the protein kinase domain of AMHR2 contains the greatest number of longevity-selected positions based on a regression analysis with 33 mammalian species ^30^. This domain contains 3 of 8 and 2 of 3 positively selected sites on branch 2 (NMR) and branch 11 (BMR), respectively.

### Different studies on positive selection in mole-rats show minor overlaps

First, we compared our list of PSGs with the PSGs detected in previous studies of positive selection in mole-rats ^18,20,21,23^ (Table S16). As observed before, ^21^ PSGs from different studies show no or small overlaps. This is not surprising because the branches examined in previous studies represent different phylogenetic entities than those used here, even though some of them are named similarly. For example, Kim et al. examined an “NMR branch” using the house mouse as closest related species ^18^In our study, the sister taxon to NMR is represented by other African mole-rats and the house mouse is used only as an outgroup (Fig. 1). In a similar way, the analysis of the African mole-rat ancestor by previous studies ^21,23^ differs from ours as we incorporated the greater cane rat as closest related short-lived species and used guinea pig as an outgroup. We therefore analyzed evolutionary processes on a shorter phylogenetic distance that closely matches the appearance of the phenotypes under investigation. In addition, there are methodological differences between the studies, e.g. regarding ortholog prediction or alignment filtering. Unfortunately, the contribution of these technical variables to the discrepancies cannot be assessed as the alignments used for the previous studies are not available and cannot be compared with those generated and provided in our study (Supplement Data).

### Positive selection and age-related regulation are linked

Next, we analyzed the direction of the regulation of PSGs during aging to identify potential links between positive selection on the analyzed branches and genetic determinants of lifespan. In general, directionality analysis of gene regulation during aging is complicated by the fact that the direction itself is not informative, whether the respective gene function is either causing or counteracting aging. E.g. up-regulation of a causative gene may accelerate aging and shorten lifespan while adaptive up-regulation to counteract aging phenotypes may extend longevity. We recently observed that PSGs in short-lived and fast-growing killifish were significantly more often up-than down-regulated during aging ^17^. This finding is consistent with the concept of antagonistic pleiotropy^31^ suggesting that the same genes that are positively selected in short-lived species for fast growth and maturation at young age are drivers of aging at old age. The antagonistic pleiotropy hypothesis is well supported, e.g. by the fact that growth rate and lifespan are negatively correlated, both between species and within many species ^6,32^. If, however, up-regulation of PSGs in short-lived species may cause aging, we hypothesized that selection for longevity is more compatible with attenuation of gene activity – either on the level of protein function or gene regulation – since avoiding damage is easier than improving repair.

Following this hypothesis, we performed RNA-seq in liver from old vs. young males of both long-lived NMRs and short-lived rats (>21 vs. 2-4 years and 24 vs. 6 months, respectively; Table S17-S19).Indeed, the union of PSGs showed preference for down-regulation in NMR and for up-regulation in rats in respect to all regulated genes (p=0.0089, Lancaster procedure ^33^). Moreover, the down-regulation in the long-lived and up-regulation in the short-lived species originate largely from the same genes as in a combined view on aging related expression changes in NMR and rat (Fig. 2), PSGs showed a highly significant preference for quadrant I (down in NMR, up in rat; p=0.0014, one-sided fisher test, quadrant I against the sum of II, III, IV). These results indicate that identified PSGs are associated with expression changes during aging of long- and short-lived rodents consistent with the antagonistic pleiotropy theory of aging.

**Figure 2.**
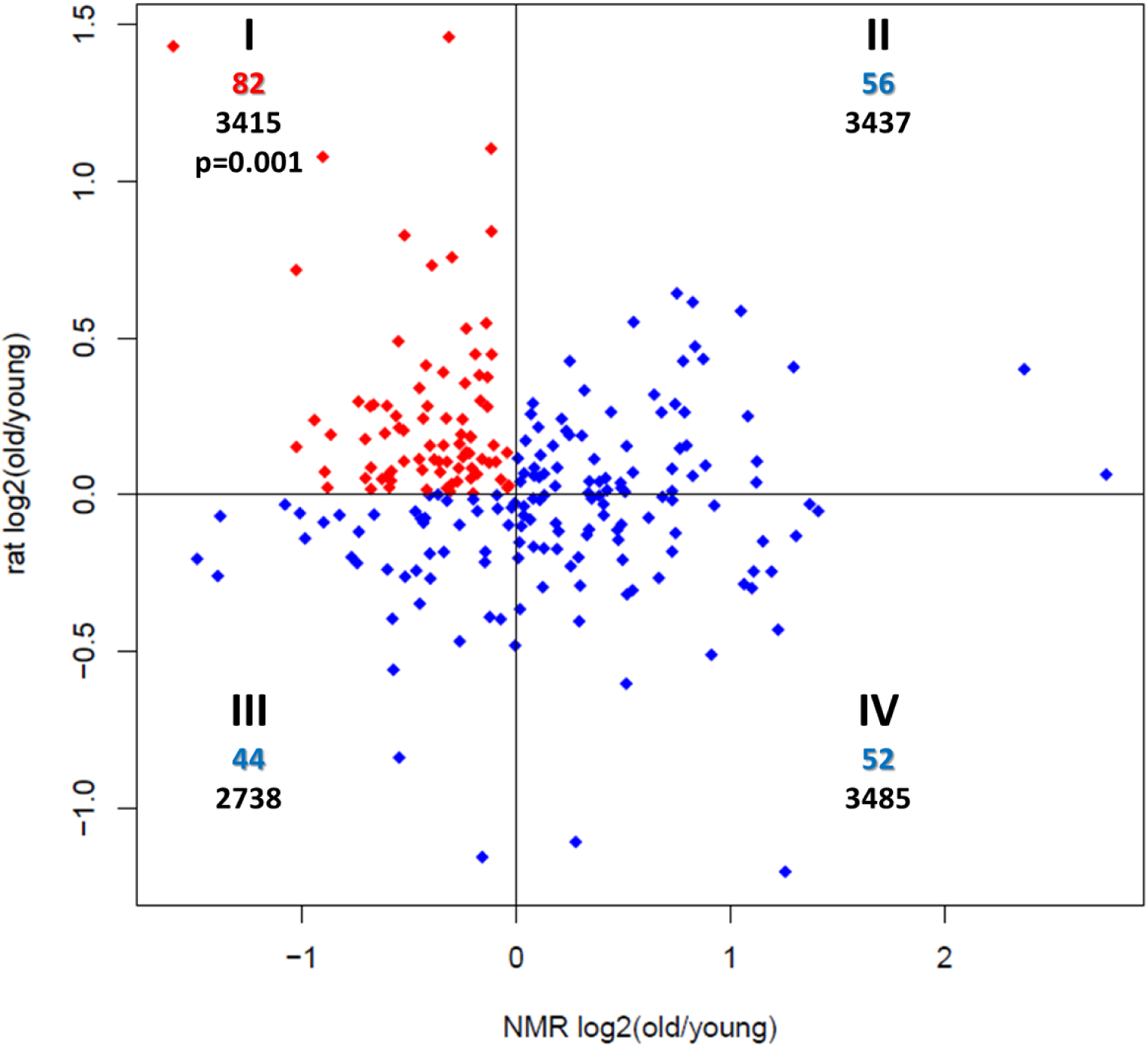
PSG expression changes during aging of NMR and laboratory rat. The roman numbers describe the quadrant, the colored numbers below that show the number of PSGs in the respective quadrant and the black numbers at the bottom give the total regulated genes in the quadrant. The red marked quadrant (I) represents PSGs that were down regulated in the long-lived NMR and up regulated in the short-lived rat and tested against the sum of three blue marked quadrants (II, III, IV) with Fisher’s exact test (one-sided). The resulting p-value is shown in quadrant I. The total number of PSGs shown in this plot (234) is lower than the unique number of all PSGs (319) due to missing expression of genes in NMR and/or rat as well as missing log2-fold-changes in at least one of the species (DEseq2).

To functionally annotate PSGs in respect to aging, we performed gene ontology (GO) term enrichment analysis. Regarding all genes, there was a significant enrichment for down-regulation in 126 terms during NMR aging while no term was enriched for up-regulation (Table S20, FDR<0.05, GAGE). The enriched 126 terms were summarized into 16 categories (Tables S21/S22, REVIGO). Among the six top categories are “translation” (GO:0006412), “cellular respiration” (GO:0045333), “response to oxidative stress” (GO:0006979) and “iron ion homeostasis” (GO:0055072) previously linked to aging (see below). With respect to possible pleiotropic effects, translation and cellular respiration are also key components of the growth program. To evaluate the PSGs in respect to these categories, we built the union of genes for each category and tested for overrepresentation of PSGs. Regarding all PSGs, there was a significant overlap with “cellular respiration” (p=0.0022, one-sided fisher test) and “response to oxidative stress” (p=0.029). Regarding only the 82 PSGs that were down-regulated in NMR and up-regulated during rat aging (quadrant I, Fig. 2), all four categories were significantly enriched (cellular respiration: p=2.1*10^-6^, response to oxidative stress: p=0.022, iron ion homeostasis: p=8.5*10^-4^, translation: p=0.011; Table S23). This again suggests that PSGs are linked to aging relevant processes in an antagonistic pleiotropic way. The result is also consistent with the hyperfunction theory of aging that suggests that antagonistic pleiotropy works via a mechanism of “perverted” growth. According to this theory the growth program that is beneficial during youth is not entirely stopped after finishing development and causes damage from that point on. The theory further claims that the master regulator mTOR governs this growth program ^34,35^

### Inflammation and host defense are enriched in branches leading to longevity

Subsequently, we searched for enriched gene ontologies in the union of PSGs across the 11 branches along which longevity evolved and in each of these branches separately (Table S24). We found enrichments of genes involved in inflammatory response (GO:0006954; FDR=0.0068, Fisher’s exact test) and defense response (GO:0006952, FDR=0.0092). Aging is tightly associated to the delicate balance between pro-inflammatory responses to resist potentially fatal infections and the inexorable damages that are accumulated by this ^36,37^. Chronic inflammation is described as a major risk factor for aging and aging-related diseases such as atherosclerosis, diabetes, Alzheimer’s disease, sarcopenia and cancer ^38^.

### mTOR, autophagy and translation pathways show signs of positive selection leading to longevity

On branch 2 (NMR), we found *RHEB* (Ras homolog enriched in brain) coding for a direct regulator of mTOR (mechanistic target of rapamycin) and on branch 9 (AMR) its paralog *RHEBL1* to be positively selected, a situation consistent with the concepts of parallel evolution as well as of subfunctionalization of genes after duplication. mTOR operates as a central regulator of cell metabolism, growth, inflammation and proliferation and was identified as a key regulator of aging and aging-related diseases in yeast, nematodes, fruit flies, and mice ^39,40^.

mTOR is also a key regulator of autophagy ^41^. Autophagy is a cellular protective cleaning mechanism, required for organelle homeostasis, especially mitochondria. While enhanced autophagy was shown to be associated with lifespan extension in worms, flies and mice, inhibition of autophagy, conversely, leads to premature aging in mice ^42^. An essential autophagy gene, *LAMP2* (lysosomal associated membrane protein 2), was identified as PSG on branch 2 (NMR) and branch 11 (BMR). As a receptor for chaperone-mediated autophagy and a major protein component of the lysosomal membrane, LAMP2 is required for degradation of individual proteins through direct import into the lysosomal lumen ^43,44^. Aging-dependent decrease of *LAMP2* expression was observed in mouse liver. Reinstatement of juvenile LAMP2 levels in aged mice significantly reduces aging-dependent decline of cell function and restores the degree of cell damage to that found in young mice ^45^.

Besides the lysosome, another cellular protein quality control and degradation system is the proteasome. While impaired proteasome function and subsequent accumulation of misfolded proteins were tightly correlated with aging and aging-related neurodegenerative disorders like Parkinson’s and Alzheimer’s disease, long-lived humans have sustained proteasome activity ^46-48^. Two proteasome subunit genes, *PSMG1* (proteasome assembly chaperone 1) and *PSMB4* (proteasome subunit beta 4), were identified as PSGs on branch 11 (BMR). PSMB4 has been classified as a driver for several types of tumors ^49^ and is a known interaction partner of PRP19 (pre-mRNA-processing factor 19 or senescence evasion factor) that is essential for cell survival and DNA repair ^50^.

Another aging relevant downstream process regulated by mTOR is translation. We identified two ribosomal proteins, RPL7L1 and RPL27A, on branch 3 (LCA of all African mole-rats except NMR). While in general, cytosolic ribosomal proteins are up-regulated with aging in humans ^51^, rats ^52^ and killifish ^53^ both genes are significantly down-regulated during NMR aging (FDR≤0.05, DESeq2). This fits the down-regulation of translation-related processes during NMR aging in general (see above). Furthermore, the protein synthesis machinery is a driver of replicative senescence in yeast ^54^. The longitudinal aging study in killifish ^55^ highlighted the starting values at 10 weeks and the amplitude of age-dependent increase of ribosomal proteins to be negatively correlated with lifespan. Inhibition of protein synthesis by reduction of ribosomal proteins was shown to extend lifespan in worms ^56^ and mice ^57^.

### Positive selection leading to longevity affects mitochondrial biogenesis and regulation of oxidative stress

Besides regulation of cytoplasmic translation of nuclear encoded genes, mTOR is also involved in mitochondrial translation. There appears to be a complex interplay between mTOR signaling, mitochondrial gene expression and oxygen consumption as well as production of reactive oxygen species (ROS) ^58-60^. Across multiple longevity-associated branches we identified PSGs that are involved in mitochondrial biogenesis (Table 1). We found, e.g., an enrichment of “mitochondrial translation” (GO:0032543, FDR=0.044) on branch 5 (SMR), the mitochondrial transcriptional termination factor (MTERF) on branch 2 (NMR) and six mitochondrial ribosomal proteins (MRPs) distributed on branches 5 (SMR), 7 (LCA of AMR and GMR) and 11 (BMR). Furthermore, nuclear encoded genes of respiratory chain complex I (*NDUFA9* and *NDUFB11*: NADH ubiquinone oxidoreductase subunits A9 and B11) and complex IV (*COX14*: cytochrome c oxidase assembly factor COX14) were identified as PSGs. Of note, we found 6 of these 15 genes to be significantly down-regulated during aging in NMR (Table 1). This suggests a functional relation of these genes to the aging process in an extremely long-lived rodent and is concordant with the down-regulation of genes involved in cellular respiration during NMR aging described above.

**Table 1.** Mitochondrial biogenesis genes under positive selection on longevity-associated branches.

| Gene | Positively selected on branch |  | NMR aging | Function in mitochondrion |
| --- | --- | --- | --- | --- |
|  | Branch number | Branch description |  |  |
| *NDUFA9* | 8 | GMR | ↓ | Complex I |
| *MTERF* | 2 | NMR | ns | Transcription |
| *SDHAF2* | 2 | NMR | ns | Complex I |
| *UQCRC2* | 2<br>3 | NMR<br>LCA after NMR<br>divergence | ↓ | Complex III |
| *NDUFB11* | 5 | SMR | ↓ | Complex I |
| *MRPS11* | 5 | SMR | ↓ | Translation |
| *MRPS36* | 5 | SMR | ↓ | Translation |
| *MRPL30* | 5 | SMR | ↓ | Translation |
| *TRNT1* | 11 | BMR | ns | RNA processing |
| *COX14* | 11 | BMR | ns | Complex IV |
| *DARS2* | 11 | BMR | ns | RNA processing |
| *MRPL57* | 11 | BMR | ns | Translation |
| *MRPL15* | 11 | BMR | ns | Translation |
| *MRPL28* | 7 | Fukomys LCA | ns | Translation |
| *GATC* | 10 | long-tailed chinchilla | ns | RNA processing |

Note: Branch numbers refer to figure 1. LCA – last common ancestor; ↓ – significantly lower expressed in liver of old animals compared to young animals (FDR<0.05); ns – not significantly changed.

Studies in mouse and the short-lived killifish have shown that expression of MRPs and complex I genes is negatively correlated with individual lifespan ^55,61^. Knock-down of MRPs in worms results in an impaired assembly of respiratory complexes and life-extension ^62^. Furthermore, we recently identified a significant enrichment of mitochondrial biogenesis genes including those for multiple MRPs, complex I components and MTERF among PSGs on two ancestral branches of annual killifishes on which lifespan was shortened considerably and independently from each other ^17^. Altogether, these results raise again the intriguing possibility that similar or even the same genes could be causally linked to the evolution of both short and long lifespan.

Mitochondria are also the main source of ROS that cause oxidative stress, i.e. damages to DNA, proteins and other cellular components ^63^. Oxidative stress is thought to play a major role in the pathogenesis of neurodegenerative diseases ^64^ and even the determination of lifespan in general (“oxidative stress theory of aging”) ^65^. On branch 3 (LCA of all African mole-rats except NMR), we found an enrichment of oxidoreductase activity (GO: GO:0016491; FDR=0.024) and positive selection of *TXN* (thioredoxin), coding for an oxidoreductase enzyme that acts as an antioxidant extending lifespan in fly ^66^ and potentially also in mice ^67,68^. As an example of continued evolution, *TXN* was found to be positively selected also on branch 7 (LCA of AMR and GMR). *SOD2* (superoxide dismutase 2) and *CCS* (copper chaperone for superoxide dismutase) are PSGs on branch 10 (chinchilla) and branch 2 (NMR), respectively. Both genes are involved in ROS defense and affect aging/lifespan in several species ^69,70^. This is interesting because in recent years, it has been repeatedly questioned that the oxidative stress theory of aging has much relevance for bathyergid rodents, given that several studies failed to find improved antioxidant capacities and/or less accumulation of oxidative damage in NMRs compared to the much shorter-lived mice ^71-73^. This is consistent with our finding of down-regulation of processes involved in “response to oxidative stress” (GO:0006979) during aging in NMR (see above). On the other hand, significantly higher levels of oxidative damage on proteins and lipids in non-reproductive as compared to reproductive females of the Damaraland mole-rat were found ^74^. Since non-reproductive individuals live shorter (and hence age faster) than their reproductive counterparts in *Fukomys* sp. ^75-77^, these results are consistent with the oxidative stress theory of aging. The diverse signs of positive selection on branch 2 (NMR), 3 (LCA of all African molerats except NMR) and 7 (LCA of AMR and GMR) may suggest that the impact of oxidative stress on aging differs between NMR and other African mole-rats.

ROS production and ROS-induced damage to biomolecules are intertwined with the formation of advanced glycation end-products (AGEs). AGEs are stable bonds between carbohydrates and proteins/lipids which are formed in a non-enzymatic fashion. AGEs activate membrane-bound or soluble AGER (AGE specific receptor) and AGEs/AGER have been linked to several aging-related diseases including Alzheimer’s disease and diabetes ^78^. Interestingly, *AGER* was found to be a PSG on branch 9 (AMR) and branch 10 (chinchilla). The role of AGEs/AGER in aging is complex and Janus-faced ^79^. AGER is significantly up-regulated in liver during NMR aging. Similarly, in skin AGE levels rise with chronological age in AMR, but surprisingly are higher in the skin of slow aging breeders than of faster aging non-breeders ^80^

### Additional links between positive selection and longevity

The gene *APOA1* (apolipoprotein A1) was found as PSG on branch 7 (LCA of AMR and GMR) and significantly up-regulated during NMR aging in liver. APOA1 is a component of HDL-particles which are as transporter of cholesterol relevant for aging-associated diseases. Polymorphisms of APOA1 are associated to coronary artery disease ^81^. Furthermore, APOA1 is an interaction partner of APOE a well-described genetic risk factor for Alzheimer’s and cardiovascular diseases ^82^ and the locus with the largest statistical support for an association with extreme longevity ^83^. Same as *MTERF* (see above), *APOA1* is one of nine genes that we recently found to be positively selected on both of two ancestral sister branches of annual fishes on which lifespan was independently reduced ^17^.

*TF* (transferrin) was identified as PSG on branch 4 (LCA of Cape, Cape dune, giant, AMR and common mole-rats). TF is an iron-binding protein responsible for transport of iron in the bloodstream and therefore essential for iron homeostasis ^84^. Neurons regulate iron intake via the TF receptor and dysregulation of this tightly controlled process in the brain is associated with neurodegenerative, age-related diseases like Parkinson’s and Alzheimer’s ^85^. TF is significantly down-regulated during NMR aging which is consistent with the down-regulation of “iron ion homeostasis” (GO:0055072) related processes during NMR aging in general (see above).

### Selection signatures of social evolution among African mole rats is consistent with a scenario of ancestral eusociality

Although all African mole-rats are strictly subterranean and occupy similar nutritional niches, intra-familiar variety of social and mating systems is amazingly high. Solitariness and polygamy in some genera (*Heliophobius*, *Georychus* and *Bathyergus*) contrast sharply with social organization in others (*Heterocephalus*, *Fukomys* and *Cryptomys*). In the latter, stable monogamous bonding of (typically one) reproductive founder pair coupled with prolonged philopatry and reproductive altruism of their offspring result in extended and cooperatively breeding family units, which can grow to considerable size. There has been much debate whether eusociality in African mole-rats is a derived or ancestral trait ^86,87^. The first scenario assumes a solitary LCA of African mole-rats (branch 1) and subsequent independent, parallel evolution of eusocial habits on branch 2 (NMR) and branch 6 (LCA of AMR, GMR and common mole-rats) and/or branch 7 (LCA of AMR and GMR) ^21^. In contrast, the second scenario suggests a eusocial LCA of all mole-rats (branch 1) and independent loss of this phenotype in the SMR (branch 5) and the LCA of the genera *Bathyergus* and *Georychus* (Cape and Cape dune mole-rat). Recent phylogenetic approaches are more supportive of scenario 2 ^88,89^ however, a PSG-based analysis of the issue is still lacking. Accordingly, we searched our data in support of one or the other scenario.

All four PSGs found on the ancestral branch 1 of all African mole-rats are involved in signaling, resulting in enrichments, e.g., of “positive regulation of cell communication” (GO:0010647, FDR=0.0092) and “positive regulation of signaling” (GO:0023056, FDR=0.0092). Enrichments in signal transduction were identified as major common pattern of the multiple independent occurrences of eusociality in bees ^90^ strongly suggesting parallel evolution in eusocial insects and mammals.

On the other hand, members of the major histocompatibility complex (MHC) were identified as PSG on branch 2 (NMR) and on branch 7 (LCA of AMR and GMR). MHC genes have a central function in the acquired immune system and immune dysfunction is involved in many neurodevelopmental disorders as well as social behavior deficits in mice and humans ^91,92^. MHCs have been implicated in language impairment and schizophrenia ^93,94^. A human allele of *HLA-A* was associated with autism defined as a pattern of behavior identified by deficits in communication and reciprocal social interactions ^95^.

Additionally, we found two signals with a potential link to social evolution on branch 2 (NMR), but not branch 6 (LCA of AMR, GMR and common mole-rats) or 7 (LCA of AMR and GMR). The first were innate immune system related enrichments “positive regulation of T cell activation” (GO:0050870, FDR=0.027) and “positive regulation of leukocyte cell-cell adhesion” (GO:1903039, FDR=0.027). In social ants and bees, it was shown that several innate immune genes have a pattern of accelerated amino acid evolution compared both to non-immunity genes in the same species and immune genes in solitary fly ^96^. Second, we found *NOTCH2* positively selected only on branch 2 (NMR). The encoded protein is one of four Notch-receptors. Notch signaling regulates interactions between physically adjacent cells and has a central role in the development of many tissues, including neurons ^97^. It was demonstrated that Notch signaling represses reproduction in worker honeybees depending on the presence of the queen and that chemical inhibition of Notch signaling can overcome the repressive effect of queen pheromone in regard to the worker ovary activity ^98^.

On the other hand, *HSD11B1* (hydroxysteroid 11-beta dehydrogenase 1) was identified as PSG on branch 6 (LCA of AMR, GMR and common mole-rats), but not branch 2 (NMR). The encoded protein catalyzes reversibly the conversion of the stress hormone cortisol to the inactive metabolite cortisone ^99^. Cortisol concentration was shown to inversely correlate with social ranks in NMRs ^100^ and with anti-social and isolation behavior in human adolescents ^101,102^. Furthermore, cortisol regulates carbohydrate metabolism that is another common enriched GO-Term in the evolution of eusociality in bees ^90^. Not linked to eusociality but still noteworthy, *HSD11B1* is significantly down-regulated during aging in the NMR and knockout of *HSD11B1* in mice improves their cognitive performance in aging ^103^. Furthermore, inhibition was described as a risk factor for cardiovascular disease and diabetes type 2 ^104^.

Taken together, our data are on best agreement with a scenario assuming eusociality (or a predisposition for it) in the LCA of all African mole-rats, followed by further independent, branch-specific evolution or loss of the phenotype leading to the distinct social genera that live today.

### Homology modeling suggests functional consequences of amino acid changes under positive selection

To evaluate the structural impact of positively selected amino acid changes, we performed homology modeling using exemplary the sites of highest probability for selection in cytoplasmic thioredoxin (TXN) and transferrin (TF). As mentioned above *TXN* is positively selected on branch 7 (LCA of AMR and GMR). In TXN of these species, there is a tyrosine residue that replaces Cys69. The latter, together with Cys62 and Cys73, constitute highly conserved mammalian non-catalytic cysteines. The local structure around Cys69 and Cys62 in TXN is important for interaction with the cytoplasmic thioredoxin reductase (TR1; ^105^), which ‘recycles’, i.e. re-reduces, the catalytic cysteines of oxidized TXN. The modeling using the structure of a fully reduced human TXN (1ERT; ^106^) as template suggests that the rather bulky side chain of Tyr69 can be accommodated in the structure of TXN (Fig. 3A), hence allowing for a productive helical interface region to TR1. TXN recycling is inhibited by formation of a disulfide bridge between Cys62 and Cys69 (Fig S1; see ^107^), e.g. under highly oxidative conditions, thereby diminishing the pool of catalytically active TXN under oxidative stress ^108,109^. Obviously, that disulfide bridge cannot form in AMR and GMR because of Cys69Tyr. From this, we conclude that Tyr69 is compatible with TXN recycling also under oxidative stress. Moreover, Cys69 is known to be a target for posttranslational modifications with impact on e.g. anti-apoptotic/apoptotic signaling pathways (for a review see: ^110^), raising interesting questions on physiological consequences of the Cys69Tyr replacement.

**Figure 3.**
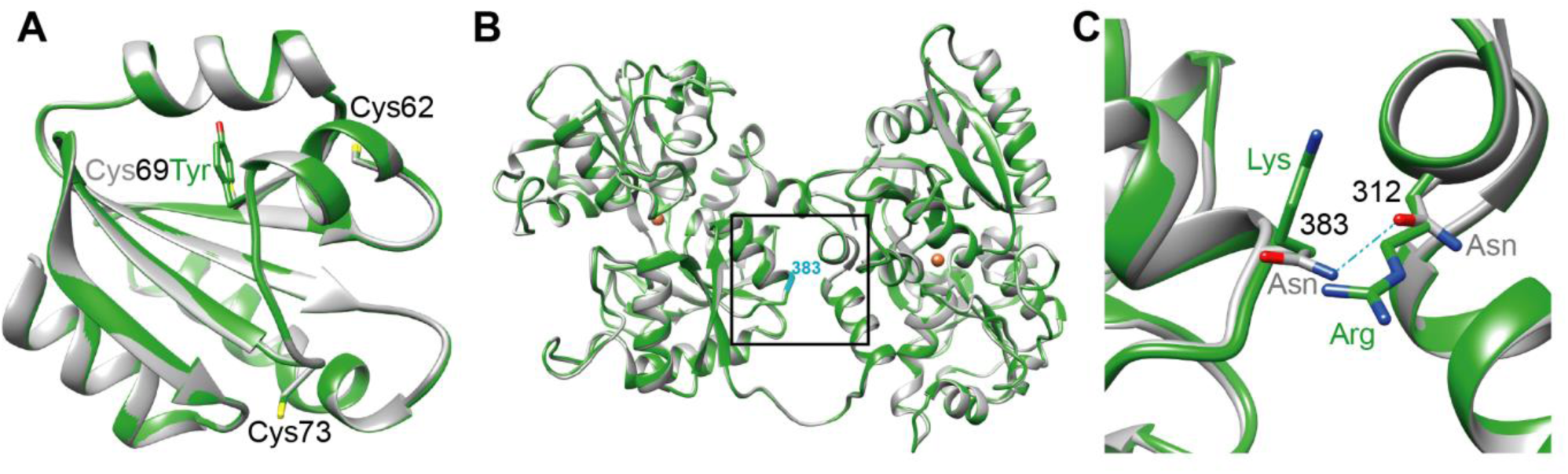
Homology models of Ansell’s mole-rat (AMR) thioredoxin (TXN) and transferrin (TF). (A) Overview of the modeled AMR TXN structure (green) superimposed onto the fully reduced human TXN template structure (1ERT, grey). Residues discussed in the text are labeled, numbering according to position in the human sequence. Color code of Cys69 and Tyr69 corresponds to the respective ribbon representation. Heteroatoms: sulfur in yellow, oxygen in red. (B) Overview of the modeled AMR TF structure (green) superimposed onto the rabbit TF template structure (1JNF, grey). The position of the Asn383Lys site discussed in the text at the boxed center of the lobe interface numbered and indicated in cyan. Brown spheres: Fe^3+^ coordinated in the template structure (1JNF, ion radius enlarged for better visibility). (C) Detail of the TF lobe interface. Shown is a magnification of the boxed region in (B). Coloring and numbering as in (B), side chain nitrogen atoms (blue), oxygen atoms (red). Potential hydrogen bond in 1JNF (light blue) as discussed in the text. Numbering (black) according to positions in the rabbit TF structure (1JNF).

TF is a PSG on branch 4 (LCA of Cape, Cape dune, giant, AMR and common mole-rats) and Ser383Lys is the site of highest probability for selection. Serum TFs form a bilobal structure, and each lobe contains two dissimilar domains with a single iron-binding site. Inspecting the structure of the AMR TF modeled on the rabbit protein (1JNF; ^111^) as template, we realized that Lys 383 is located at the interface between the two lobes (Fig. 3B). In the rabbit TF two juxtapositioned Asn residues at position 383 and 312 might form an H-bond and this constellation could stabilize the inter-lobe interactions (Fig. 3C). In contrast, the juxtaposition of the positively charged side chains of Lys383 and a conserved Arg312 in the AMR structural model (Fig. 3C) would be expected to weaken the lobe-lobe interaction due to electrostatic repulsion. The functional consequences for TF or TXN implied by this modeling require experimental validation.

## Conclusions

We provided a systematic scan for PSGs on evolutionary branches of the African mole-rat family and other rodents leading to longevity and eusociality. Due to the incorporation of species from all six genera of the African mole-rats as well as its closest relative, the cane rat, into the analysis, we were able to examine considerably more extant and ancestral branches than previous studies. This enabled the analysis to provide a high resolution of positive selection on branches on which the mentioned traits had evolved.

Analyzing the gene expression of PSGs, we found a highly significant pattern of down-regulation in the long-lived NMR and up-regulation in the short-lived rat, fitting the antagonistic pleiotropy theory of aging ^112^ and the hyperfunction theory of aging. The latter claims mTOR as a central hub affecting aging and lifespan ^35^. Correspondingly, the PSGs and enriched functional terms cover many of the processes that are regulated by the mTOR pathway, e.g. translation, autophagy and mitochondrial biogenesis. Furthermore, with *RHEB* and its ortholog *RHEB1L* direct regulators of mTOR ^113^ are under positive selection in two of the branches. In addition, we linked positive selection with immune system and the antioxidant defense, processes known to be involved in regulation of lifespan.

With regard to evolution of eusociality, our findings are in line with the theory that the LCA of all African mole-rats had at least a predisposition for social lifestyle that was lost in some lineages, while in other lineages the ancestral phenotype has further evolved, leading to the distinct social phenotypes in the extant species.

Moreover, we exemplarily showed potential functional relevance of the positively selected sites by homology modeling on the protein level. This may encourage experimental follow-up studies since all sequences and alignments including the identified positively selected sites are accessible via supplement data.

## Methods

### CDS data

We examined nine African mole-rat species covering all six genera. Additionally, our analysis comprises eight outgroup species, including the long-lived BMR and the chinchilla. mRNA sequences of seven distantly related outgroup species were obtained from RefSeq along with their CDS annotation (Table S1). For the NMR we used a recently published *de novo* transcriptome assembly ^114^. RNA-seq data for six mole-rat species was obtained from GenBank Sequence Read Archive, study SRP061925 ^21^. The reads were assembled and annotated using FRAMA as described in ^114^.

For AMR and GMR, purification of RNA from 13 and 17 tissues, respectively, was done using Qiagen RNeasy Mini Kit following the manufacturer’s description. Novel RNA-seq was performed for both species as described in Table S2. *De novo* transcriptome assemblies of the generated data were performed using FRAMA ^114^ (see Table S1).

For the SMR and the greater cane rat genome sequencing was performed to complement the transcriptome data. DNA was isolated from liver tissue of two female SMR individuals and a male liver of the greater cane rat using DNeasy Blood & Tissue (Qiagen). DNA was then converted to Illumina libraries and sequencing was done as given in Table S2. Sequence reads were cleaned by removal of adaptors and low-quality regions at the ends (i.e. regions with more than 10% with quality score ≤ 20). Low quality reads (i.e. less than 50% remained) and duplicons were discarded. *De novo* genome sequence assembly was performed using CLC assembler (CLC Genomics) with default settings. The CDS annotation was done using AUGUSTUS ^115^ with AMR CDSs as hint.

All animals were housed and euthanized compliant with national and state regulations. Read data was deposited as ENA (European Nucleotide Archive) study PRJEB20584.

### Identification of positively selected genes

To scan on a genome-wide scale for genes under positive selection, we fed the CDSs of the described species set along with the branches we wanted to examine (Fig. 1) into the PosiGene pipeline ^28^.GMR was used as PosiGene’s anchor species. Regarding the SMR, for which we had both a genome and a transcriptome assembly, we used generally the transcriptome assembly, except for those ortholog groups in which no SMR ortholog could be assigned via transcriptomic but via genomic data. This was accomplished by calling the three PosiGene modules separately, feeding both assemblies independently in the first module (ortholog assignment) and deleting all genome-based SMR sequence in those ortholog groups that contained transcriptome-based SMR CDSs before calling the second module. An overview about the number of genes and sequences tested for positive selection in the different branches is shown in Table S1. We considered all genes with nominal p-values ≤ 0.05 as PSGs.

### Gene ontologies

We determined enrichments for GO categories with Fisher’s exact test based on on the R package GOstats. The resulting p-values were corrected using the Benjamini-Hochberg method (FDR).

### Differentially expressed genes during NMR and rat aging

The young and old rats (strain Wistar) had an age of 6 (n=4) and 24 (n=5) months, respectively. The young NMRs had an age of 3.42±0.58 years (average±sd, n=6). The old NMRs were at least 21 years old (recorded lifetime in captivity, n=3). All examined animals were males. All animals were housed and euthanized compliant with national and state regulations. For both species, purification of RNA from liver samples was done using Qiagen RNeasy Mini Kit following the manufacturer’s description. In short, we performed RNA-seq using Illumina HiSeq 2500 w ith 50 nt single read technology and a sequencing depth of at least 20 mio reads/sample (Table S17). For NMR, the read mapping was performed with STAR ^116^ (--outFilterMismatchNoverLmax 0.06 --outFilterMatchNminOverLread 0.9 -- outFilterMultimapNmax 1) against the public genome (Bioproject: PRJNA72441) that we had annotated before by aligning the above mentioned NMR transcriptome reference using BLAT ^117^ and SPLIGN ^118^. Rat reads were aligned against the mentioned RefSeq reference using bwa aln ^119^ (-n 2 -o 0 -e 0 -O 1000 -E 1000). Read data and counts were deposited as GEO (Gene Expression Omnibus) series GSE98746. Differentially expressed genes (FDR≤0.05, Table S18, S19) and fold-changes were determined with DESeq2 ^120^. GAGE ^121^ was used to determine enriched gene ontologies based on fold-changes (Table S20). Gene ontologies with FDR≤0.05 were summarized using REVIGO (allowed similarity=0.5) ^122^. Four of the six largest summarized categories of the resulting treemap (Table S21/S22) were further analyzed due their aging relevance (representative terms given): “translation” (GO:0006412), “cellular respiration” (GO:0045333), “response to oxidative stress” (GO:0006979) and “iron ion homeostasis” (GO:0055072). For each of these categories the union of genes across gene ontology terms was built. These unions were tested for significant overlaps with (i) the union of PSGs across branches and (ii) the union of PSGs across branches that were down-regulated during aging in NMR and up-regulated in rat (Fisher’s exact test). Functional annotation of the PSGs in respect to the four categories is given in Table S23).

### Homology modeling of protein structure

Models were built in SWISS-MODEL (http://swissmodel.expasy.org;) ^123,124^. No further optimization was applied to the resulting TXN and TF models. Superimposition of the model and template structures and rendering was carried out using CHIMERA ^125^.

### Data availability

Read data for AMR, GMR, SMR and greater cane rat was deposited as ENA (European Nucleotide Archive) study PRJEB20584. Read data for NMR and rat was deposited as GEO (Gene Expression Omnibus) series GSE98746.

## Acknowledgements

We thank Ivonne Görlich, Christiane Vole and Yoshiyuki Henning for excellent assistance, Debra Weih for proofreading the manuscript and Christoph Kaether for helpful discussions. This work was funded by the Deutsche Forschungsgemeinschaft (DFG, PL 173/8-1 and DA 992/3-1), the European Community’s Seventh Framework Programme (FP7-HEALTH-2012-279281) as well as the Leibniz association (SAW-2012-FLI-2).

## Author contributions

MP, PD and KS initiated the project. MP, AS and KS managed the project. PD, HB, TH, SH and MS provided samples; MGr was in charge for the sequencing; MB and AS performed assemblies and gene expression analyses; AS searched for PSGs; AS and HAK applied statistical tests; MGö performed protein structure modeling; AS, MP, PD, AC, HB, CC and CM interpreted the data; AS, MP, PD, AC and MGö wrote the manuscript. All authors read and approved the final manuscript.

## Corresponding author

Correspondence to Arne Sahm (http://arne.sahm@fli-leibniz.de).

## Supplementary information

**Supplementary tables: S1-S24.xls**

Table S1. Data Sources for assemblies and sequence statistics.

Table S2. Samples that were sequenced to create genome/transcriptome assemblies.

Table S3. PSGs on multiple branches.

Table S4. Overview of positively selected genes (FDR≤0.05) on examined branches.

Table S5. Results on branch 1.

Table S6. Results on branch 2.

Table S7. Results on branch 3.

Table S8. Results on branch 4.

Table S9. Results on branch 5.

Table S10. Results on branch 6.

Table S11. Results on branch 7.

Table S12. Results on branch 8.

Table S13. Results on branch 9.

Table S14. Results on branch 10

Table S15. Results on branch 11.

Table S16. Overlaps between this and previos studies.

Table S17.Samples that were RNA-sequenced to examine gene regulation during aging.

Table S18. DESeq2 result for gene expression comparison of young (Ø 3.42 years) vs. old (>21 years) naked mole-rats.

Table S19. DESeq2 result for gene expression comparison of young (6 months) vs. old (24 months) rats.

Table S20. GAGE gene ontology enrichment for expression changes during NMR aging (Ø 3.42 vs > 21 years, FDR<=0.05, all down-regulated)

Table S21. REVIGO treemap result of GAGE enrichment for differential expression during NMR aging.

Table S22. REVIGO representative categories (representative term given) of GAGE enrichment for differential expression during NMR aging.

Table S23. PSGs in aging relevant summarized REVIGO categories and quadrant 1 (up-regulated in rat and down-regulated in NMR).

Table S24. Gene ontologies enriched for PSGs on examined branches based on GOStats and Fisher’s exact test (FDR≤0.05).

**Supplement data:** ftp://genome.leibniz-fli.de/pub/mrps2017/supplement_data.tar.gz

The package contains visualizations of alignments and positively selected sites for all genes and branches that were analyzed in this article.

